# Tracing a protein’s folding pathway over evolutionary time using ancestral sequence reconstruction and hydrogen exchange

**DOI:** 10.1101/334607

**Authors:** Shion A. Lim, Eric R. Bolin, Susan Marqusee

**Affiliations:** Department of Molecular and Cell Biology, University of California, Berkeley, Berkeley, CA, United States; Institute for Quantitative Biosciences (QB3), University of California, Berkeley, Berkeley, CA, United States; Biophysics Graduate Program, University of California, Berkeley, Berkeley, CA, United States; Department of Chemistry, University of California, Berkeley, Berkeley, CA, United States; The authors contributed equally to this work

**Keywords:** protein folding, hydrogen exchange, protein evolution, ancestral sequence reconstruction, mass spectrometry

## Abstract

The conformations populated during protein folding have been studied for decades; yet, their evolutionary importance remains largely unexplored. Ancestral sequence reconstruction allows access to proteins across evolutionary time, and new methods such as pulsed-labeling hydrogen exchange coupled with mass spectrometry allow determination of folding intermediate structures at near amino-acid resolution. Here, we combine these techniques to monitor the folding of the ribonuclease H family along the evolutionary lineages of *T. thermophilus* and *E. coli* RNase H. All homologs and ancestral proteins studied populate a similar folding intermediate despite being separated by billions of years of evolution. Even though this conformation is conserved, the pathway leading to it has diverged over evolutionary time, and rational mutations can alter this trajectory. Our results demonstrate that evolutionary processes can affect the energy landscape to preserve or alter specific features of a protein’s folding pathway.

## Introduction

Protein folding, the process by which an unfolded polypeptide chain navigates its energy landscape to achieve its native structure,^1,2^ can be defined by the partially folded conformations (intermediates) populated during this process. Such intermediates are key features of the landscape; they can facilitate folding, but they can also lead to misfolding and aggregation, resulting in a breakdown of proteostasis and disease.^3–5^ While identifying and characterizing these intermediates is critical to understanding and engineering a protein’s energy landscape, their transient nature and low populations present experimental challenges. Recent technological improvements in hydrogen exchange monitored by mass spectrometry (HX-MS) have provided access to the structural and temporal details of these folding intermediates at near-single amino-acid resolution.^6–11^ This pulsed-labeling HX-MS approach is particularly well suited to studies of multiple variants or families of proteins, as it does not require large amounts of purified protein or NMR assignments. Thus, pulsed-labeling HX-MS can be used to address long-standing questions in the field: How robust is a protein’s energy landscape to changes in the amino acid sequence, and how conserved is the folding trajectory over evolutionary time?

Ribonuclease HI (RNase H) is an ideal system to investigate protein folding over evolutionary time. RNase H from *E. coli*, ecRNH* (the asterisk denotes a cysteine-free variant of RNase H), is arguably one of the best-characterized proteins in terms of its folding pathway and energy landscape. Both stopped-flow ensemble studies and single-molecule optical trap experiments demonstrate that this protein populates a major obligate intermediate before the rate-limiting step in folding.^12–16^ A rare population of this intermediate can also be detected under native-state conditions.^17^ Several homologs of RNase H have also been studied, yielding insight into the folding trends of extant RNases H.^18–20^

In addition to comparing the folding pathways of homologs, one can use a phylogenetic technique called ancestral sequence reconstruction (ASR) to access the evolutionary history of a protein family and study the properties of ancestral proteins.^21,22^ ASR has been applied to a variety of protein families and in addition to revealing the evolutionary history, these ancestral proteins can act as intermediates in sequence space to uncover mechanisms underlying protein properties.^23–30^ Recently, ancestral sequence reconstruction was applied to the RNase H family and the thermodynamic and kinetic properties of seven ancestral proteins connecting the lineages of *E. coli* and *T. thermophilus* RNase H (ecRNH* and ttRNH*) were characterized.^31–33^ Stopped-flow kinetics monitored by circular dichroism (CD) demonstrate that all seven ancestral proteins populate a folding intermediate before the rate-limiting step. Additionally, the folding and unfolding rates show notable trends along the phylogenetic lineages, and the presence of a folding intermediate plays an important role in modulating these evolutionary trends.^32^

For ecRNH*, multiple methods have confirmed the structural details of the folding intermediates. This major folding intermediate, termed I_core_, which forms before the rate limiting step, involves secondary structure from the core region of the protein, including Helices A-D and Strands 4 and 5, while the rest of the protein (Helix E and Strands 1, 2, 3), remains unfolded (Figure 1A).^12,34,35^ Pulsed-labeling HX-MS with near amino acid resolution was developed using ecRNH* as the model protein.^6^ This approach confirmed the structure of I_core_ and revealed the stepwise protection of individual helices leading up to the intermediate. Specifically, the amide hydrogens in Helix A and Strand 4 are the first elements to gain protection, followed by those in Helix D and Strand 5, and then Helices B and C to form the canonical I_core_ intermediate. The periphery, comprising of Strands 1-3 and Helix E, gains protection in the rate-limiting step to the native state. Would this I_core_ folding intermediate and the stepwise folding pathway be conserved across evolution?

**Figure 1.**
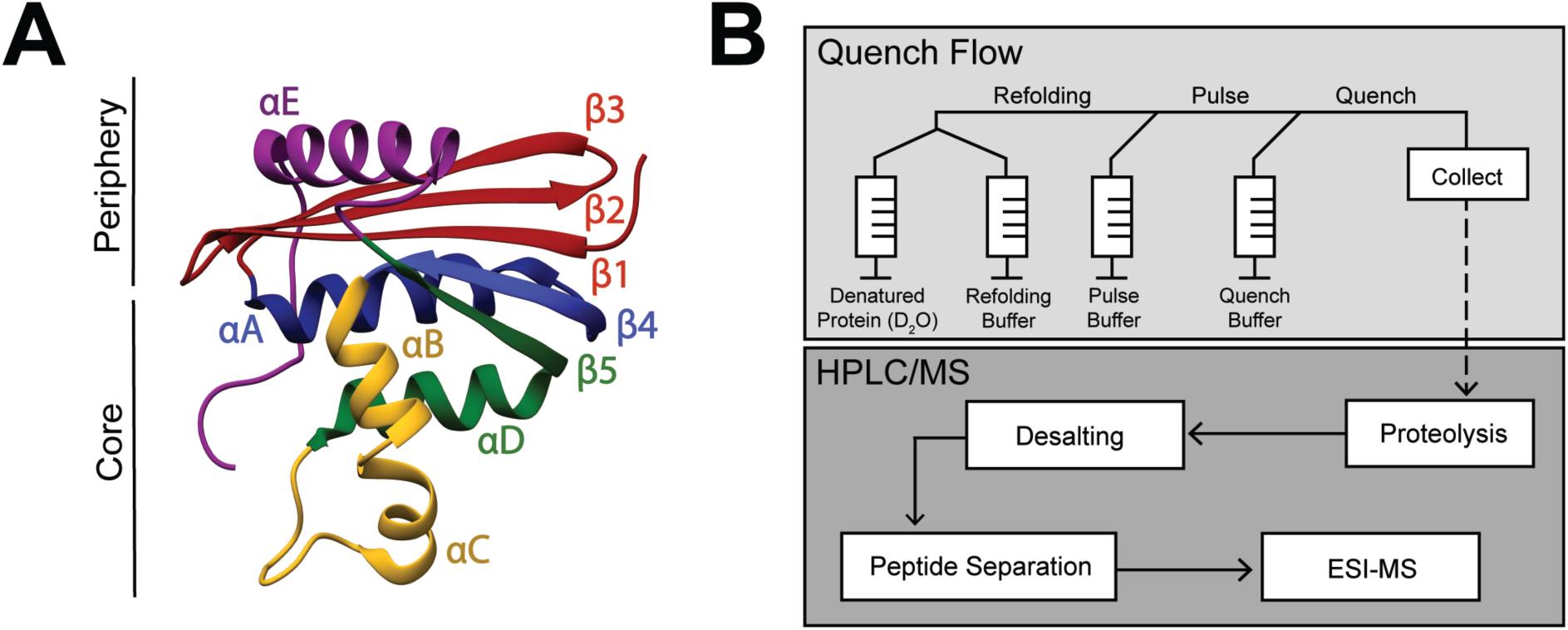
RNase H structure and Pulsed-labeling HX-MS. **A)** Crystal structure of *E. coli* RNase H* (ecRNH*) (PDB: 2RN2).^54^ Secondary structural elements: Red: Strand 1, Strand 2, Strand 3 (S123); Blue: Helix A, Strand 4 (HAS4); Yellow: Helix B, Helix C (HBHC); Green: Helix D, Strand 5 (HDS5); Purple: Helix E (HE). The core region of the protein (I_core_) involving Helix A, Strand 4, Helix B, Helix C, Helix D, Strand 5 and the periphery region of the protein involving Strand 1, Strand 2, Strand 3, Helix E are denoted. **B)** Pulsed-labeling setup and workflow. Unfolded, fully deuterated protein in high [urea] is rapidly mixed with low [urea] refolding buffer to initiate refolding. After some refolding time, hydrogen exchange of unprotected amides is initiated by mixing with high-pH pulse buffer. The hydrogen exchange reaction is quenched by mixing with a low-pH quench buffer. The sample is injected onto an LC-MS for in-line proteolysis, desalting, and peptide separation by reverse-phase chromatography followed by MS analysis.

Here, we use pulsed-labeling HX-MS on the resurrected family of RNases H to investigate the evolutionary and sequence determinants governing the folding trajectory. Specifically, we find that the structure of the major folding intermediate (I_core_) has been conserved over three billion years of evolution, suggesting that this partially folded state plays a crucial role in the folding or function of the protein. The detailed steps leading to this folding intermediate, however, vary. The very first step in folding differs between the two extant homologs: for ecRNH*, Helix A gains protection before Helix D, while for ttRNH*, Helix D acquires protection before Helix A. This pattern can be followed along the evolutionary lineages: most of the ancestors fold like ttRNH* (Helix D before Helix A) and a switch to fold like ecRNH* (Helix A before Helix D) occurs late along the mesophilic lineage. These phylogenetic trends allow us to investigate how these early folding events are encoded in the amino acid sequence. By selectively modulating biophysical properties, notably intrinsic helicity, of specific secondary structure elements, we are able to favor or disfavor the formation of specific conformations during folding and have engineering control over the folding pathway of RNase H.

## Results

### Monitoring a protein’s folding trajectory by pulsed-labeling HX-MS

We used pulsed-labeling hydrogen exchange monitored by mass spectrometry (HX-MS) on extant, ancestral, and site-directed variants of RNase H to examine the robustness of a protein’s folding pathway to sequence changes. These experiments allow us to characterize the partially folded intermediates and the order of structure formation during folding to ask whether these intermediates have changed over evolutionary time, and what role sequence might play in determining these intermediates.

Figure 1B outlines the scheme for the pulsed-labeling experiment (for details, see Methods). Briefly, folding is initiated by rapidly diluting an unfolded (high [urea]), fully deuterated protein into folding conditions (low [urea]) at 10°C. After various folding times (t_f_), a pulse of hydrogen exchange is applied to label amides in regions that have not yet folded. The amount of exchange at each folding timepoint is then detected by in-line proteolysis and LC/MS. Data are analyzed first at the peptide level by monitoring the protection of deuterons on peptides as a function of refolding time, and then at the residue level, using overlapping peptides de-convoluted by the program HDsite.^36,37^

Since the original folding studies on RNase H were carried out at 25°C, we re-characterized the folding of each RNase H variant at 10°C using stopped-flow circular dichroism spectroscopy (Figure S1). The refolding profiles were consistent with those at 25°C.^12,19,32^ At low [urea], all ancestors show a large signal change (burst phase) within the dead time of the stopped-flow instrument (∼15 msec), followed by a slower observable phase which fit well to a single exponential. The resulting chevron plots (ln(*k*_obs_) vs [urea]) show the classic rollover at low [urea] due to the presence of a stable folding intermediate. As expected, the observed rates at 10°C are slower than 25°C, but the chevron profiles are similar for all RNase H variants. Thus the overall folding trajectory, notably the population of a folding intermediate, has not changed between the two temperatures.

### Monitoring the folding pathway of ttRNH* using pulsed-labeling HX-MS

First, we characterized the conformations populated during folding of extant RNase H from *T. thermophilus* and compared its folding trajectory to the previously characterized folding trajectory of *E. coli* RNase H.^6^ 374 unique peptides were identified by MS. Of these, 49 unique peptides were observed at all refolding time points and were used for further analysis (Figure 2A). Similar to ecRNH*, peptides associated with I_core_ (Helix A-D, Strands 4-5) gain protection early (within milliseconds), corresponding to the timescale for the formation of the folding intermediate. Peptides associated with the periphery of the protein (Strands 2-3, Helix E) gain protection on the order of seconds, corresponding to the rate-limiting step (Figure 2B). Thus, the major folding intermediate in ttRNH*, I_core_, is strikingly similar to that of ecRNH*.^6^

**Figure 2.**
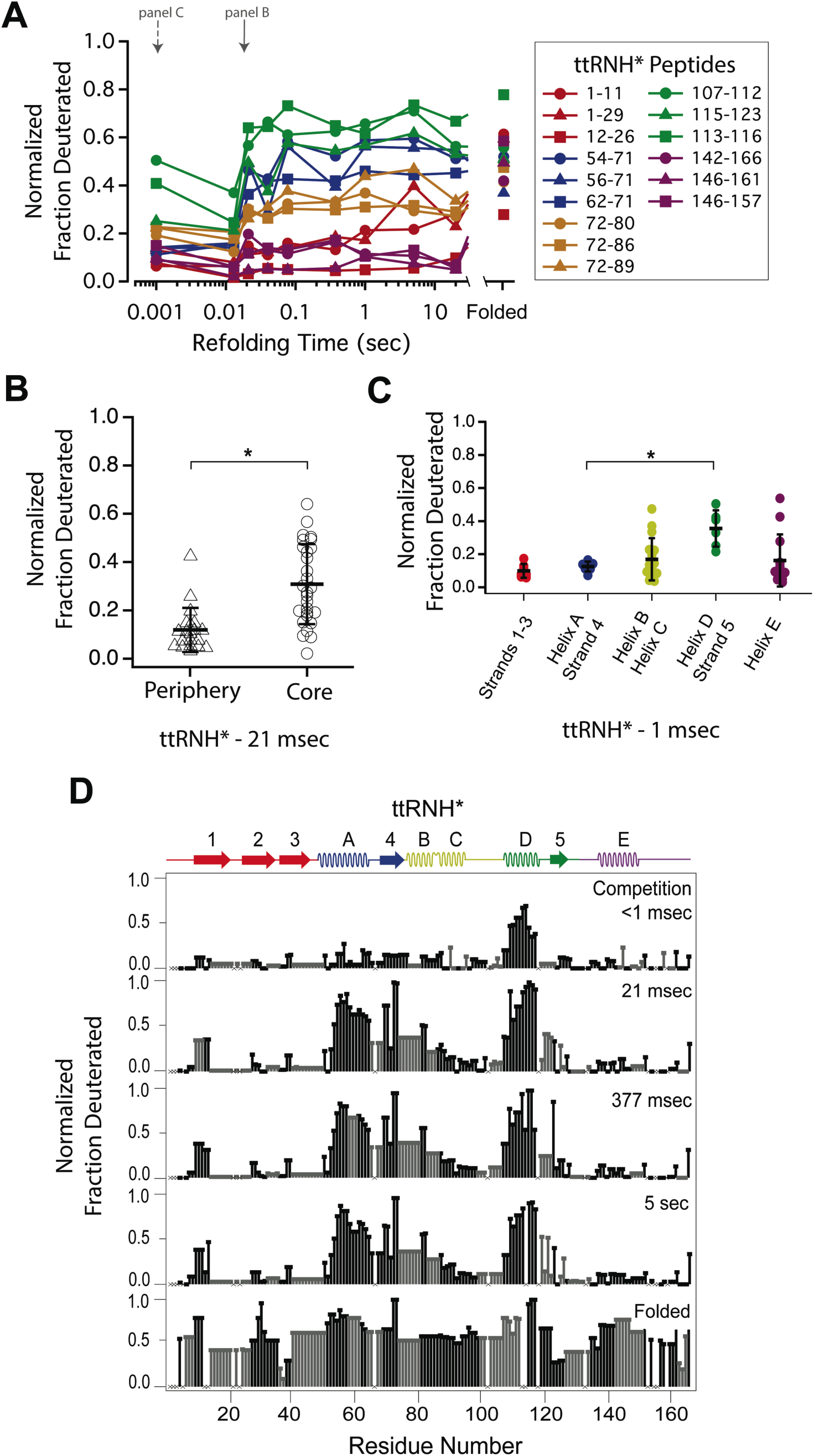
Determination of the folding pathway of *T. thermophilus* RNase H* by HX-MS. **A)** Protection of representative peptides from ttRNH* at various refolding times. Peptides are colored according to their corresponding structural element. The solid arrow indicates the refolding time point analyzed in panel B. The dotted arrow indicates the refolding time point analyzed in panel C. **B)** Protection of peptides mapping to the core region (I_core_) or the periphery region of ttRNH* at 21 msec after refolding. Bars represent the mean and standard deviation of each data set. *p < 0.0001 (Welch’s unpaired T-test) **C)** Protection of peptides of ttRNH* mapping to distinct secondary structural elements at 1 msec after refolding. Bars represent the mean and standard deviation of each data set. *p = 0.0027 (Welch’s unpaired T-test). **D)** Residue-resolved folding pathway of ttRNH* at representative refolding time points. Data points in black indicate residues that are site-resolved. Data points in grey indicate residues in regions with less peptide coverage and are thus not site-resolved with the neighboring residues. Residues where site-resolved protection could not be determined due to insufficient peptide coverage is denoted with a “x”.

Looking at the very early refolding times allows one to determine the individual folding steps preceding I_core_. At the earliest time point (∼1 msec), almost all peptides are unfolded (fully exchange with solvent) with the exception of those in Helix D and Strand 5, which are ∼40% deuterated (Figure 2C). Peptides spanning Helix A and Strand 4 are less protected (∼15% deuterated) at this same time point. This order of protection (Helix D before Helix A) is notably different than that for *E. coli* RNase H*, where Helix A is protected before Helix D.^6^ Peptides spanning Helix B and Helix C gain protection in the I_core_ intermediate. Peptides from Strands 1-3 and Helix E do not gain full protection until significantly later (on the order of seconds), corresponding to the rate-limiting step to the native state. Thus, while the I_core_ intermediate is largely conserved between ttRNH* and ecRNH*, the initial steps of folding differ between the two homologs.

The peptide data from each time point were also analyzed using HDSite to determine residue-level protection in a near site-resolved manner (Figure 2D). These site-resolved data also show protection appearing first in Helix D and Strand 5, followed by Helix A/Strand 4, Helix B/C, and finally, the periphery Helix E and Strands 1-3. The differences in the order of protection leading up to I_core_ of ecRNH* and ttRNH* are also evident in this site-resolved analysis.

### Pulsed-labeling HX-MS on the ancestral RNases H

To look for evolutionary trends in the folding trajectory, we probed the folding pathway of ancestral RNases H along the lineages of *E. coli* and *T. thermophilus* RNase H (Figure 3A). Anc1* is the last common ancestor of ecRNH* and ttRNH*. Anc2* and Anc3* are ancestors along the thermophilic lineage leading to ttRNH*, and AncA*, AncB*, AncC*, and AncD* are ancestors along the mesophilic lineage leading to ecRNH*. Previous kinetic studies demonstrated that all of the ancestral proteins fold via a three-state pathway, populating an intermediate before the rate-limiting step.^32,33^ We now use pulsed-labeling HX-MS to obtain a near-site resolved trajectory of the folding pathway for each ancestor and determine whether the I_core_ structure is conserved over evolution.

**Figure 3.**
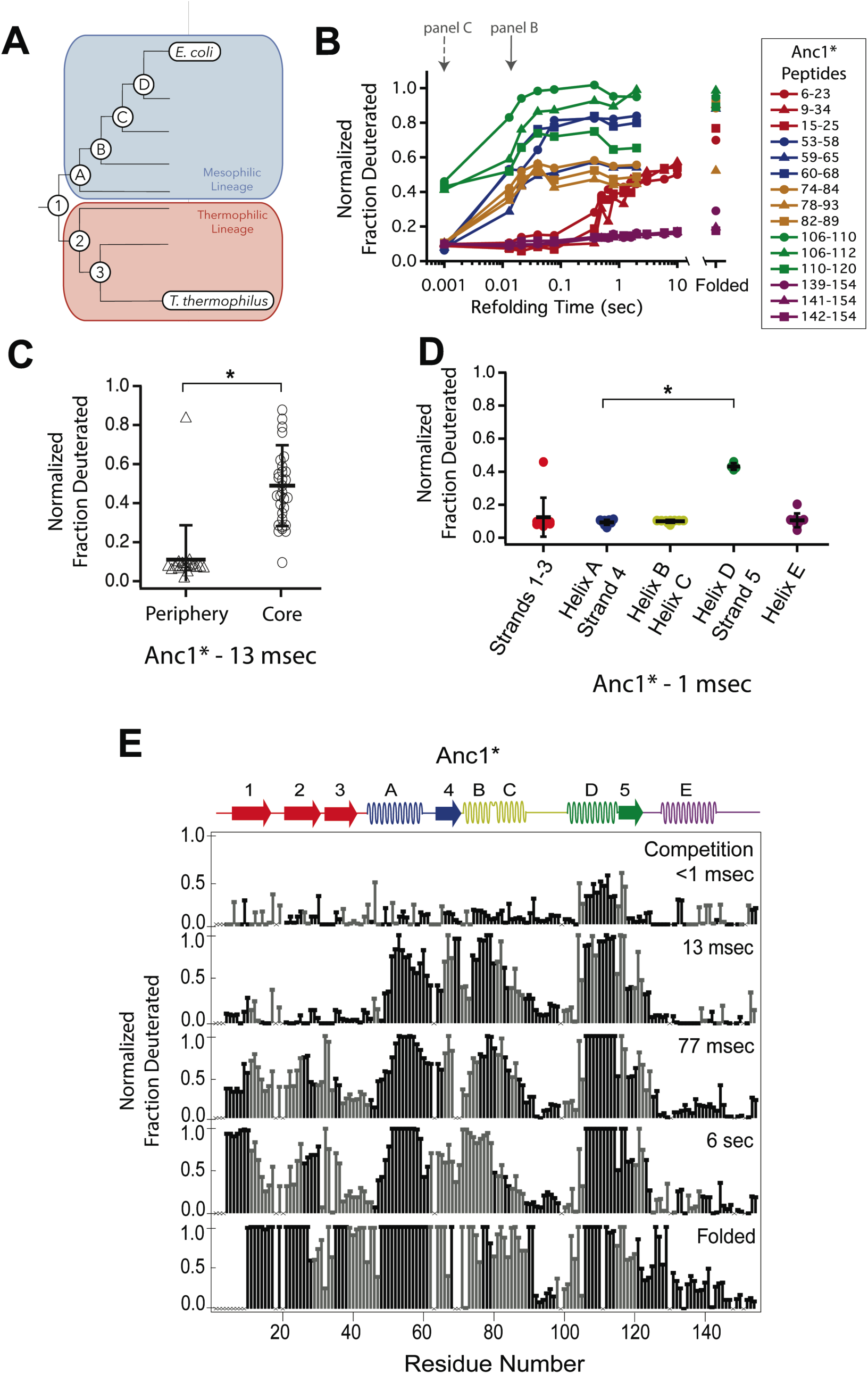
Determination of the folding pathway of ancestral RNases H by HX-MS. **A)** Representation of the phylogenetic tree of the RNase H family illustrating the ancestral proteins along the two lineages leading to *E. coli* RNase H and *T. thermophilus* RNase H. Adapted from Figure 2A of Hart KM et al. 2014, *PLoS Biology*. 12(11) doi:10.1371/journal.pbio.1001994, published under the CreativeCommons Attribution 4.0 International Public License (CC BY 4.0; https://creativecommons.org/licenses/by/4.0/).^31^ Anc1* is the last common ancestor of ecRNH* and ttRNH*. Anc2* and Anc3* are ancestors along the thermophilic lineage to ttRNH*. AncA*, AncB*, AncC*, and AncD* are ancestors along the mesophilic lineage to ecRNH*. **B)** Protection of representative peptides from Anc1* at various refolding times. Peptides are colored according to their corresponding structural element. The solid arrow indicates the refolding time point analyzed in panel B. The dotted arrow indicates the refolding time point analyzed in panel C. **C)** Protection of peptides mapping to the core region (I_core_) or the periphery region of Anc1* at 13 msec after refolding. Bars represent the mean and standard deviation of each data set. *p = 0.0011 (Welch’s unpaired T-test) **D)** Protection of peptides mapping to distinct secondary structural elements of Anc1* at 1 milliseconds after refolding. Bars represent the mean and standard deviation of each data set. *p < 0.0001 (Welch’s unpaired T-test). **E)** Residue-resolved folding pathway of Anc1* at representative refolding time points. Data points in black indicate residues that are site-resolved. Data points in grey indicate residues in regions with less peptide coverage and are thus not site-resolved with the neighboring residues. Residues where site-resolved protection could not be determined due to insufficient peptide coverage is denoted with a “x”.

We obtained good peptide coverage for all of the ancestors with a minimum of 81 peptides seen in all time points for each variant (Figure 3, Figures S2-S7). As observed in both ttRNH* (above) and ecRNH*^6^ all of the ancestral RNases H populate the canonical I_core_ folding intermediate prior to the rate-limiting step. Peptides corresponding to the I_core_ region of the RNase H structure become protected on the timescale of milliseconds, while the rest of the protein gains protection on the timescale of seconds (Figure 3C, Figures S2-S7). Thus, the structure of this major folding intermediate is not only present in both extant RNases H, but is conserved over nearly three billion years of evolutionary history.

Similarly to the extant proteins, the periphery of the ancestral proteins gains protection on a much slower timescale (Figure 3C, Figure S2-S7). The details of protection in this region, however, vary somewhat across the ancestors. The periphery becomes fully protected by the last time point in all ancestral proteins except for AncB* (Figure S5). AncB* was previously characterized to be non-two-state with a notable population of the folding intermediate under equilibrium conditions,^32^ and the lack of protection in the periphery in the folded state of AncB* is consistent with this observation. For Anc1* and Anc2*, there are also notable differences in the time course of protection for the terminal helix, Helix E. For these two proteins, the peptides spanning Helix E are decoupled from Strands 1-3 (which show protection on the same timescale as global folding) and do not gain protection even in the folded state of the protein (Figure 3B, Figure 3D, Figure S2), suggesting that Helix E is improperly docked or poorly structured in Anc1* and Anc2*. Indeed, Helix E is known to be labile in ecRNH*: a deletion variant of ecRNH* without this final helix forms a cooperatively folded protein,^38^ and recent single-molecule force spectroscopy of ecRNH* showed that Helix E can be pulled off the folded protein under low force while the remainder of the protein remains structured (manuscript in preparation). It appears that Helix E may be further destabilized in Anc1* and Anc2* such that it does not show protection in the native state.

### The early folding steps of RNase H change across evolutionary time

Since the order of events leading to I_core_ differs between the extant homologs, we examined whether the ancestral RNases H spanning the lineages of these two homologs show any trends in their early folding steps. For each ancestor, we analyzed the fraction of deuterium protected in peptides that are uniquely associated with specific helices of the protein (Figure 3D and Figure S2-S7) to determine which regions fold first.

These data show that the last common ancestor of ecRNH* and ttRNH*, Anc1*, as well as all proteins along the thermophilic lineage (Anc2* and Anc3*) show similar behavior to ttRNH* and gain protection first in Helix D/Strand 5 (Figure S2, S3). For the first two ancestors along the mesophilic lineage (AncA* and AncB*), the order of protection is difficult to determine. For AncA*, there is no significant difference in the degree of protection among the peptides within I_core_ (this analysis is limited by the availability of peptides associated exclusively within a region) (Figure S4). However, when all overlapping peptides are analyzed using HDSite to obtain site resolution, we observe notable protection in Helix D at the earliest refolding times. Therefore, we conclude that although Helix D folding before Helix A is likely, the early folding events of AncA* cannot be unambiguously determined. For AncB*, all of I_core_ gains protection at the same time point, both at the peptide and residue-level, so the order of assembly cannot be determined with our time resolution (Figure S5).

The next ancestor along the mesophilic lineage, AncC*, shows protection first in Helix D, indicating that this pattern of protection is maintained through the mesophilic lineage to this ancestor (Figure S6). AncD*, the most recent ancestor along the mesophilic lineage, however, is similar to ecRNH* and gains protection first in Helix A (Figure S7). As detailed for the other ancestors, the data were also analyzed using HDSite to determine residue-level protection for each ancestral RNase H (Figure 3E, Figure S2-S7). These data indicate a pattern in the order of protection in the early steps of the folding pathway across the RNase H ancestors. Early protection in Helix D is an ancestral feature of RNase H that is maintained in the thermophilic lineage, with a transition occurring late during the mesophilic lineage to a different pathway where Helix A is protected before Helix D, resulting in a distinct folding pathway for the two extant RNase H homologs (Figure 4).

**Figure 4.**
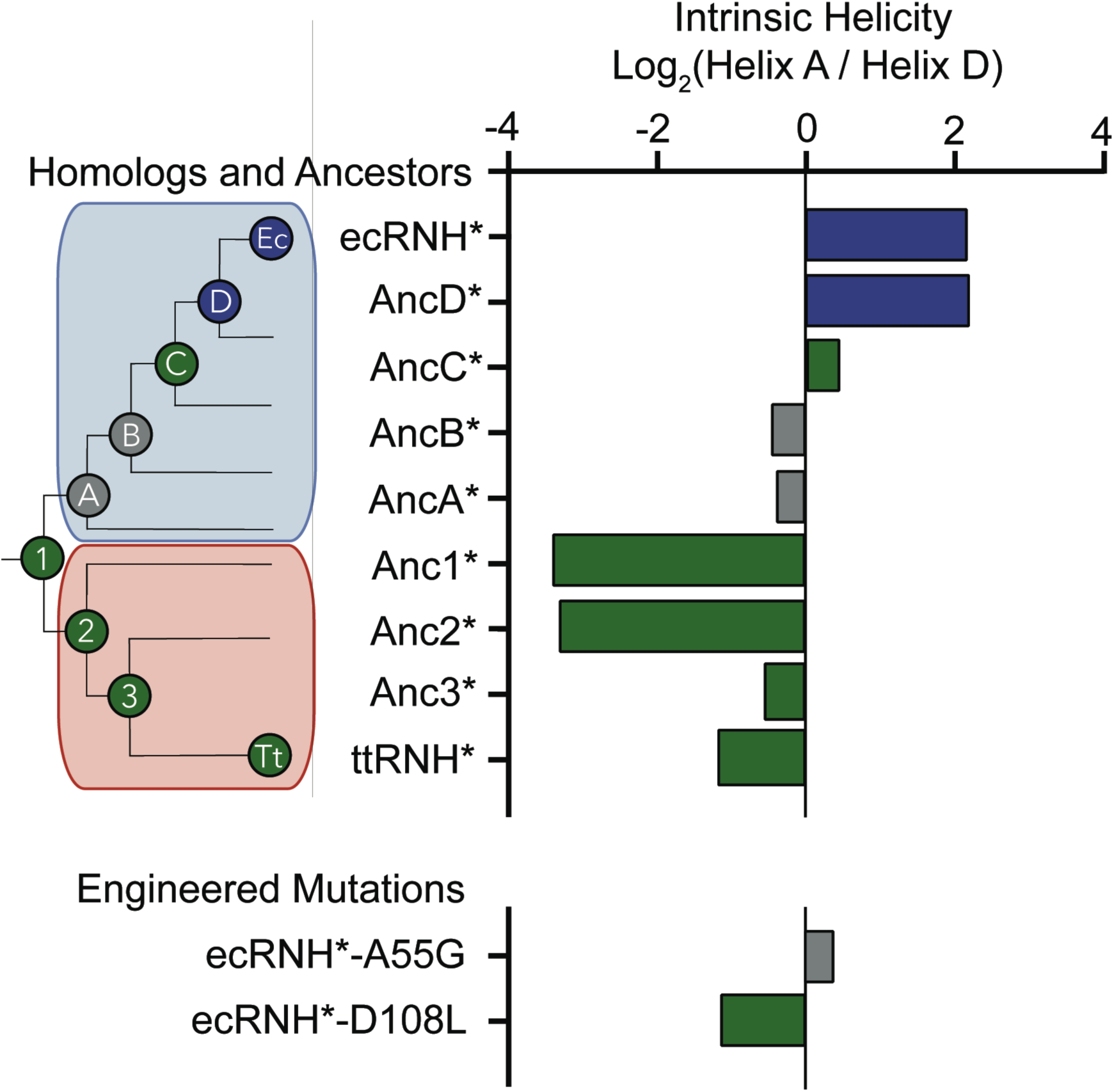
Intrinsic helicity as a predictor for the early folding mechanism of RNases H. Log-ratio of intrinsic helicity of Helix A and Helix D for each RNase H variant studied. Intrinsic helix predictions were calculated using AGADIR.^39^ The order of helix protection for each variant of RNase H is depicted in color. Green bars represent proteins where Helix D is the first structural element to gain protection during refolding. Blue bars represent proteins where Helix A is the first structural element to gain protection during refolding. Grey bars represent proteins where the helix protection order could not be unambiguously determined. The order of helix protection for each ancestor and homolog is also colored on the phylogenetic tree, revealing a trend in the RNase H folding trajectory along the evolutionary lineages. The phylogenetic tree shown in this figure is adapted from Figure 2A of Hart KM et al. 2014, *PLoS Biology*. 12(11) doi:10.1371/journal.pbio.1001994, published under the CreativeCommons Attribution 4.0 International Public License (CC BY 4.0; https://creativecommons.org/licenses/by/4.0/).^31^

### Early helix protection is determined by the local sequence of the core

Relative to the vast sequence space available, these RNase H ancestors represent a set of closely related sequences with distinct folding properties and provide an excellent system to help us elucidate the physiochemical mechanism and the sequence determinants dictating the RNase H folding trajectory. An analysis of the intrinsic helical propensity of each region using the algorithm AGADIR^39^ shows a notable trend in helicity that correlates with the early folding events (Figure 4). For proteins that gain protection in Helix A first, the intrinsic helicity of Helix A is four-fold higher than that of Helix D. For the variants where Helix D is protected first, the intrinsic helicity of Helix D is similar to or greater than Helix A. This suggests that intrinsic helix propensity may play an important role in determining which region is the first to gain protection during the folding pathway of RNase H. To investigate this hypothesis, we turned to rationally designed variants.

### Intrinsic helicity plays a role in determining the structure of the early intermediates

If the order of protection in the early folding events of RNase H is determined by intrinsic helix propensity, then we should be able to alter the protein sequence rationally and manipulate the folding trajectory. Thus, we asked whether single-site mutations that change the relative helix propensity of Helix A and Helix D could alter the folding trajectory of ecRNH* and make it fold in a similar fashion to ttRNH*. Two different point mutations were made in ecRNH*: A55G decreases helix propensity in Helix A, and D108L increases helicity in Helix D (Figure 4, Figure 5A, Table S1). Pulsed-labeling HX-MS indicates that both of these variants alter the early folding events of ecRNH*. The peptide-level protection of ecRNH* A55G indicates that at 13 msec, both Helix A and Helix D show similar levels of protection. In contrast, for wild-type ecRNH*, Helix A shows protection by 1 msec and Helix D does not show comparable protection until 10-20 msec.^6^ Thus, the mutation A55G slows the gain of protection in Helix A such that it no longer protected before Helix D (Figure 5B). The peptide-level protection of ecRNH* D108L indicates a change in the order of protection. Due to the limited number of peptides available, we could only confidently determine this using peptides spanning the N-terminus of Helix D. At 13 msec, the N-terminus of Helix D (residues 106-108) near the D108L mutation is protected significantly faster than any other region of the protein. Thus increasing helix propensity correlated with a change in the folding trajectory. (Figure 5C). Together, these two mutations suggest that intrinsic helicity plays a role in the early folding events of RNase H and can be used to alter the stepwise order of conformations populated during folding.

**Figure 5.**
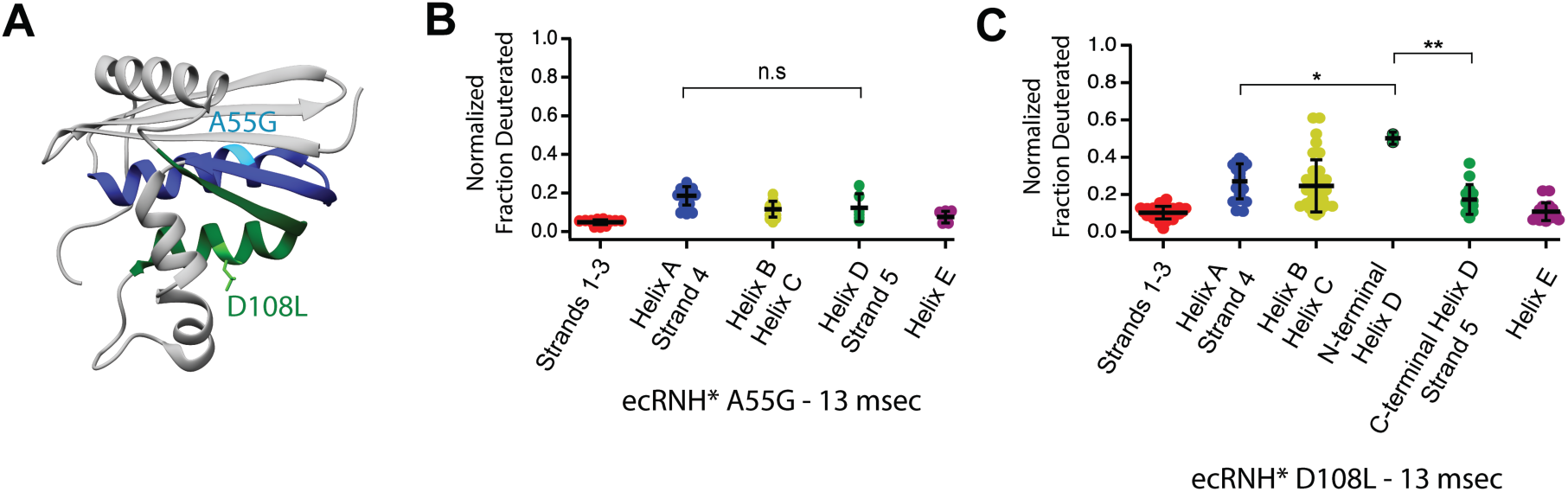
Engineered mutations to alter the folding pathway of ecRNH*. **A)** Crystal structure of *E. coli* RNase H (PDB: 2RN2) with mutations designed to alter intrinsic helicity.^55^ A55G, located in Helix A (blue), is colored in cyan. D108L, located in Helix D (green), is colored in light green. **B)** Protection of peptides mapping to distinct secondary structural elements of ecRNH* A55G at 13 msec after refolding. Bars represent the mean and standard deviation of each data set. p = 0.0917 (n.s. = not significant, Welch’s unpaired T-test). **C)** Protection of peptides mapping to distinct secondary structural elements of ecRNH* D108L at 13 msec after refolding. Bars represent the mean and standard deviation of each data set. *p = 0.0016, **p = 0.0044 (Welch’s unpaired T-test)

## Discussion

### Determining the folding pathway of multiple protein variants

Pulsed-labeling hydrogen exchange is currently the most detailed method to identify the conformations populated during protein folding. This approach was initially developed for use with NMR detection where it benefited from NMR’s site-specific resolution of individual amides.^40^ However, using NMR with pulsed-labeling HX requires tens of milligrams of sample and NMR peak assignments for the amides in each protein studied. In addition, probes are limited to amide sites stable to exchange in the final folded state (protection factors of >∼80,000) resulting in loss of information at individual sites, which can sometimes represent large regions of the protein. In contrast, detection by mass spectrometry as applied in this study requires much less protein sample, has much faster data collection, and can theoretically cover 100% of the protein sequence. Importantly, this approach does not demand any structural information of the folded state, such as NMR assignments, for the specific protein or variant studied. These advantages enabled us to obtain the stepwise folding pathway of nine variants of RNase H and study the evolutionary history and sequence determinants of the RNase H folding pathway in detail. While pulsed-labeling HX-MS has been used to characterize the folding pathways of several model systems, this study is the first to utilize the higher throughput nature of HX-MS to study an ensemble of protein variants. The advantages of this technique to study many different sequences of the same fold shows great promise for probing the relationship between amino acid sequence and a protein’s energy landscape and will likely be particularly valuable for protein engineering and design applications.

### I_core_ is a structurally conserved folding intermediate over 3 billion years of evolution

The native fold of a protein is robust to changes in sequence, proteins with >∼30% sequence identity share the same fold.^41^ Thus small variations in sequence, such as those found among homologs or site-specific mutations, do not affect the overall three-dimensional structure of a protein. These mutations can, however, affect the overall energy landscape, which in turn can have profound effects of function. Here, we find conservation of a high-energy structure populated during the folding of the RNase H family over incredibly long evolutionary timescales. Using pulsed-labeling HX-MS we identified and characterized the structure of the major folding intermediate in seven ancestral and several mutant RNases H, which together with previous studies on extant homologs, suggest that the conservation of this intermediate is a key feature of the RNase H energy landscape across ∼3 billion years of evolutionary time.

Why does I_core_ persist on the energy landscape of RNase H? One explanation is a simple topological constraint; all RNases H may need to fold via a populated I_core_ intermediate to successfully reach the native state. This explanation, however, is countered by a previous study where a single mutation (I53D) in ecRNH* destabilizes I_core_ such that it is no longer populated during folding—yet this variant still folds to the native state.^34^ Adding osmolytes, such as sodium sulfate, stabilizes this folding intermediate and switches ecRNH* I53D back to a three-state folding pathway, showing that the presence of the folding intermediate can be modulated. Additionally, a fragment of RNase H containing only the I_core_ sequence (and variants thereof) can autonomously fold and be studied at equilibrium, indicating that this structure is stable and robust to mutations.^42,43^ The nature of the rate-limiting step, or folding barrier, which allows for the buildup of this intermediate is unclear. One possibility is that the I_core_ intermediate is populated simply because the information for folding this region is completely encoded locally and I_core_ can fold relatively fast, before this rate limiting step to the fully folded state.

Alternatively, I_core_ could be conserved because it contributes to the biological function or fitness of the protein. Partially folded states and high-energy non-native conformations are known to be important for a variety of protein functions and proteostasis.^4,44,45^ All of the ancestral RNases H we studied here are active, in that they cleave RNA-DNA hybrids in vitro;^31^ and although the residues thought to contribute to substrate-binding affinity are contained in the core region of the protein,^46^ the active site residues (D10, E48, D70) span both the core and the periphery. It is therefore possible that a stable folding core with an energetically independent periphery is important for the efficiency or dynamics associated with catalysis in RNase H.

While the presence of the I_core_ intermediate has been observed in all proteins studied here, recent studies have suggested that some of the RNase H variants, notably for proteins along the thermophilic lineage, the I_core_ folding intermediate may also involve structure in the first β-strand.^33,43,47^ While we see slight protection in this region for ttRNH*, hydrogen exchange may not be the best probe of this—docking of Strand 1 without its hydrogen-bonding partners in the rest of the β-sheet may not be reflected by backbone amide protection. Therefore, amide protection may not be observed even if Strand 1 docks early to the core. The involvement of Strand 1 in ancestral other RNase H variants studied remains unclear from this study.^33,43^

### Aspects of the folding pathway are malleable across evolutionary time

Our pulsed-labeling HX-MS results also illustrate how other features of a protein’s energy landscape can be altered over evolutionary timescales. Although the I_core_ intermediate is conserved across all RNases H studied, the individual folding steps leading up to I_core_ differ. Anc1*, the last common ancestor, folds through a pathway where the Helix D/Strand 5 region is the first structural element to gain protection. This ancestral feature is maintained along the thermophilic lineage to the extant ttRNH*. Along the mesophilic branch, we observe a switch from this ancient folding pathway to one that first forms protection in Helix A/Strand 4 that occurs evolutionarily between AncC* and AncD*. This suggests that while the structure of I_core_ has been conserved across 3 billion years of evolution, the steps to form this intermediate are malleable over time. Since an isolated helix is unlikely show protection by HX, we expect additional hydrophobic collapse of the polypeptide to contribute to the observed protection. Nonetheless, the switch in protection between Helix A and Helix D indicates that formation of native structure nucleates in a different region of the protein across the RNase H variants studied, with a clear evolutionary trend.

Despite these trends, it remains difficult to rationalize these observations in terms of a selective evolutionary pressure or fitness implication. These very early events occur on the order of one millisecond, significantly faster than the overall folding of the protein. Furthermore, all of these RNase H proteins fold to their native state efficiently with no evidence for aggregation or misfolding. So, although partially folded states have been implicated as gateways for aggregation for some proteins,^4^ this does not appear to be the case for RNase H. It is possible that the change in the early folding step is a result of mutations that are coupled to another feature under selection or drift. Although the actual evolutionary implication for the RNase H folding pathway may be lost in history, the trend in folding pathway across evolutionary time demonstrates that folding pathways and conformations on the energy landscape of proteins can be affected over time, and this system provides an excellent tool to interrogate the role sequence plays in guiding the process of protein folding.

### The folding pathway of RNase H can be altered using simple sequence changes

Our study also shows how insights from evolutionary history can contribute to our understanding of the physiochemical mechanisms dictating the protein energy landscape and how we might use that knowledge to engineer the landscape. The regions that gain protection first involve helical secondary structure elements, and their folding order correlates with isolated helical propensity of these regions predicted by AGADIR.^39^ Proteins where protection is first observed in Helix A have higher intrinsic helicity in Helix A than in Helix D. Proteins where Helix D gains protection first higher helicity in Helix D or roughly equal helicity in both regions This property was used to guide our site-directed mutagenesis to select variants to alter the folding trajectory of ecRNH* in a predictive manner using intrinsic helicity as a guide.

While these results are consistent with local helicity as a determinant of the earliest folding steps, there may be other parameters that dictate the formation of these conformations. The parameter average area buried upon folding (AABUF)^48^ which measures the average change in surface area of a residue from an unfolded state to a folded state, has been shown to correlate to the structure of the folding intermediate in apomyoglobin.^49,50^ Both helicity and AABUF are altered in the mutants considered in our study (Table S1). Indeed, AABUF and helicity are often correlated and contributions of either parameter are difficult to disentangle. Nevertheless, our data suggest that parameters that are locally encoded in regions of a protein can be used engineer the energy landscape of a protein including its folding pathway.

We have used a combination of ASR and pulsed-labeling HX-MS to explore the conformations populated during the folding of multiple RNase H proteins, including homologs, ancestors, and single-site variants. All RNase H proteins studied populate the same major folding intermediate, I_core_, indicating that this conformation has been maintained on the energy landscape of RNase H over long evolutionary timescales (>3 billion years). This remarkable conservation of a partially folded structure on the energy landscape of RNase H is contrasted with changes in the folding pathway leading up to this structure. The early folding events preceding this intermediate (Helix A protected before Helix D or vice versa) differ between the two homologs and also shows a notable trend along the evolutionary lineages. This pattern of protection correlates with the relative helix propensity of the sequences comprising these two helices, and we use this knowledge to alter the folding pathway of ecRNH* through rationally designed mutations. Our study illustrates how the energy landscape of a protein can be altered in complex ways over evolutionary time scales, and how insights from evolutionary history can contribute to our understanding of the physiochemical mechanisms dictating the protein energy landscape.

## Acknowledgements

We thank members of the Marqusee Lab for helpful discussions. We also thank Leland Mayne and members of the Englander Lab and Goran Stjepanovic in the Hurley Lab for support on the pulsed-labeling HX-MS instrumental setup and analysis. We thank the Vincent J. Coates Proteomics/Mass Spectrometry Facility for instrumentation support. This work was funded by NIH Grant GM050945 (to S.M.) and a National Science Foundation Graduate Research Fellowship (to S.A.L).

## Materials and Methods

### Protein Purification

Cysteine-free *T. thermophilus* RNase H, and ancestral RNases H were expressed and purified as previously described.^31,51,52^ Point mutants were generated using site-directed mutagenesis, confirmed by Sanger sequencing, and the proteins were purified as previously described.^53^ Purity was confirmed by SDS-PAGE and mass spectrometry.

## HX-MS System

Hydrogen exchange mass spectrometry (HX-MS) experiments were carried out using a system similar to that described by Mayne et al.^7,8^ Briefly, a Bio-Logic SFM-4/Q quench flow mixer with a modified head piece with reduced swept volume was used to initiate protein refolding, followed by pulse-labeling unprotected amide hydrogen atoms, and quenching of the labeling reaction. The minimum dead time for mixing is 13 msec. Quenched samples were injected into an HPLC system constructed using two Agilent 1100 HPLC instruments. The quenched sample was flowed over columns (Upchurch C130B) packed with beads of immobilized pepsin and fungal protease at 400 µL/min in 0.05% TFA. The digested protein was run onto a C-4 trap column (Upchurch C-128 with POROS R2 beads) for desalting. An acetonitrile gradient (15-100% acetonitrile, 0.05% TFA at 17 µL/min) eluted peptides from this C-4 trap column and onto an analytical C-8 column (Thermo 72205-050565) for separation before injection into an ESI source for mass spectrometry analysis on a Thermo Scientific LTQ Orbitrap Discovery. The entire HPLC system is kept submerged in an ice bath at 0°C to reduce back exchange of deuterium atoms during the chromatography steps. The workflow takes ∼10-18 minutes from injection to peptide detection.

## Refolding Experiment

Similar to previous reports,^6,8^ unfolded protein samples in high denaturant (80 µM [protein], 20 mM NaOAc pH=4.1, 7-9 M [urea]) were deuterated by a repeated cycle of lyophilization and resuspension in D_2_O. For the pulsed labeling experiment, 1 volume of deuterated protein was mixed in the SFM-4/Q with 10 volumes of refolding buffer (10 mM Sodium Acetate pH=5.29, H_2_O) to initiate refolding. The pulse for hydrogen exchange was initiated by mixing with 5 volumes of high pH buffer (100 mM Glycine pH=10.11) and then quenched after 10 msec with 5 volumes low pH buffer (200 mM Glycine pH=1.95). The length of the delay line between the first and second mixer was changed to achieve a range of refolding times. An interrupted mixing protocol was used to measure the longest refolding time points (>373 msec). Undeuterated protein was used to perform tandem mass spectrometry (MS/MS) analysis to compile a list of peptides and their retention times in the HPLC system. Competition experiments where refolding and exchange were initiated at the same time were performed by diluting deuterated protein in high urea into high-pH refolding buffer (100 mM Glycine pH=10.11). In this experiment each site will exchange with the solvent around it unless it can gain protection before exchange occurs (<1 msec on average). For each time point, an identical sample was collected in which the high pH pulse was replaced by unbuffered water to measure back exchange for each sample. All data were obtained in triplicate and were normalized for back exchange. Data for ttRNH* were normalized to the theoretical maximum number of deuterons as back exchange controls for this protein did not produce enough peptides. Fully folded controls were created by diluting unfolded protein samples 1:10 in fully deuterated refolding buffer and incubating at room temperature for 4 hours before applying the same 10 msec high-pH pulse using the SFM-4/Q.

## MS detection and data analysis

Proteome Discoverer 2.0 (Thermo Scientific) was used to identify peptides from the tandem MS data. Peptides identified in the pulse-labeled refolding experiments with deuterated protein were used to determine the presence and deuteration level of each peptide at each refolding time point. The spectral envelope of each peptide was fit using two separate algorithms developed by the Englander Lab to determine their deuteration state — ExMS for identification and fitting of peptides and HDsite for deconvolution of overlapping peptides to achieve near-amino acid level deuteration levels.^36,37^ In addition, HDExaminer (Sierra Analytics) was used to identify and fit each peptide and determine deuteration levels. Different charge states of the same peptide were averaged where noted and used for further analysis. Centroids of each peptide at each time point taken from HDExaminer were used for further analysis. The residue cutoffs for specific structural regions of each protein were determined from a multiple sequence alignment using the structure of *E. coli* RNase H as a guide (PDB: 2RN2).^31^ Peptides were assigned to different structural regions based on these residue cutoffs. Peptides that spanned multiple secondary structural regions of a protein were excluded from further analysis, as were peptides not present in all time points. Peptides mapping to Strands 1-3 and Helix E were assigned to the periphery region of the protein. Peptides mapping to Helix A-D and Strands 4-5 were assigned to the core region of the protein.

## Supplemental Materials

Tracing a protein’s folding pathway over evolutionary time using ancestral sequence reconstruction and hydrogen exchange

**Figure S1.**
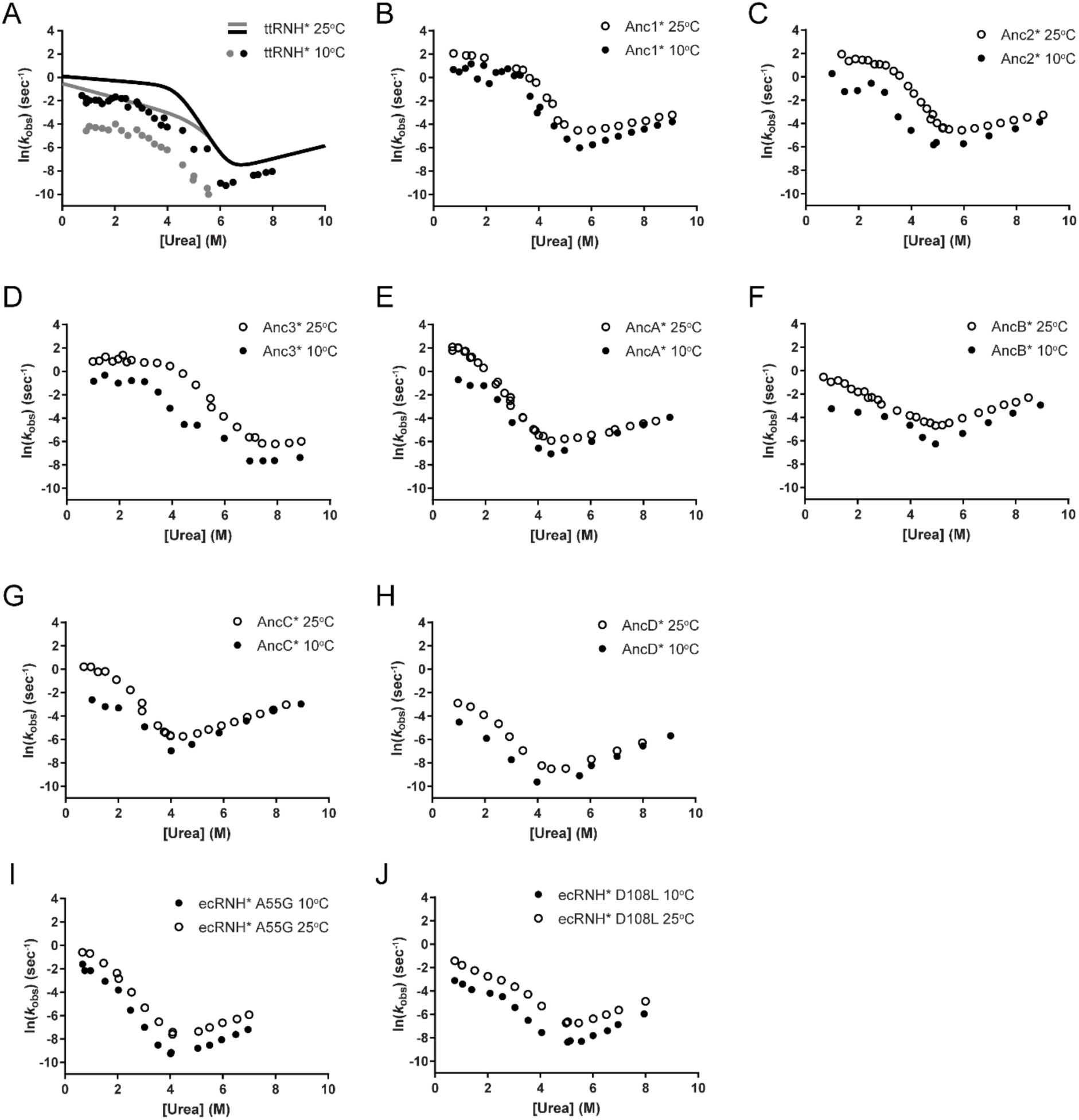
Chevron plot of RNase H variants studied at 10°C and 25°C. Chevron plots (ln(*k*_obs_) vs [urea]), determined from refolding and unfolding experiments in various [urea] at 10°C and 25°C for **A)** ttRNH*, **B)** Anc1*, **C)** Anc2*, **D)** Anc3*, **E)** AncA*, **F)** AncB*, **G)** AncC*, **H)** AncD*, **I)** ecRNH* A55G, **J)** ecRNH* D108L. For **A)** Both the fast (black dots) and slow (grey dots) rates of folding for ttRNH* are shown at 10°C, and chevron fits for the two rates at 25°C are shown as lines and adapted from previous work.^1^ Data at 25°C for **B)** – **H)** were adapted from a previously published study.^2^

**Figure S2.**
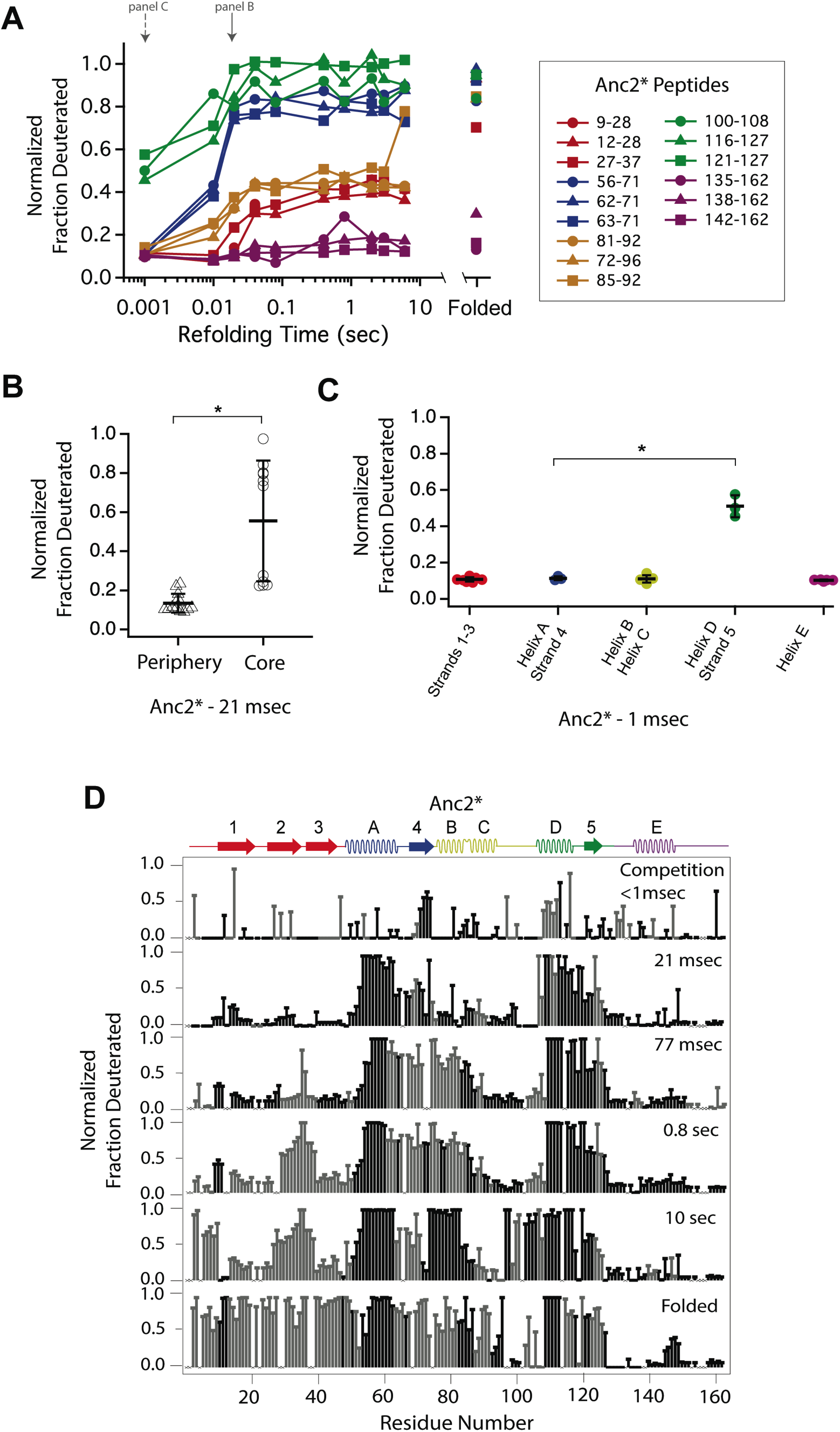
Determination of the folding pathway of Anc2* by HX-MS. **A)** Protection of representative peptides from Anc2* at various refolding times. Peptides are colored according to their corresponding structural element. The solid arrow indicates the refolding time point analyzed in panel B. The dotted arrow indicates the refolding time point analyzed in panel C. **B)** Protection of peptides mapping to the core region (I_core_) or the periphery region of Anc2* at 21 msec after refolding. Bars represent the mean and standard deviation of each data set. *p = 0.0011 (Welch’s unpaired T-test) **C)** Protection of peptides of Anc2* mapping to distinct secondary structural elements at 1 msec after refolding. Bars represent the mean and standard deviation of each data set. *p = 0.0064 (Welch’s unpaired T-test). **D)** Residue-resolved folding pathway of Anc2* at representative refolding time points. Data points in black indicate residues that are site-resolved. Data points in grey indicate residues in regions with less peptide coverage and are thus not site-resolved with the neighboring residues. Residues where site-resolved protection could not be determined due to insufficient peptide coverage is denoted with a “x”.

**Figure S3.**
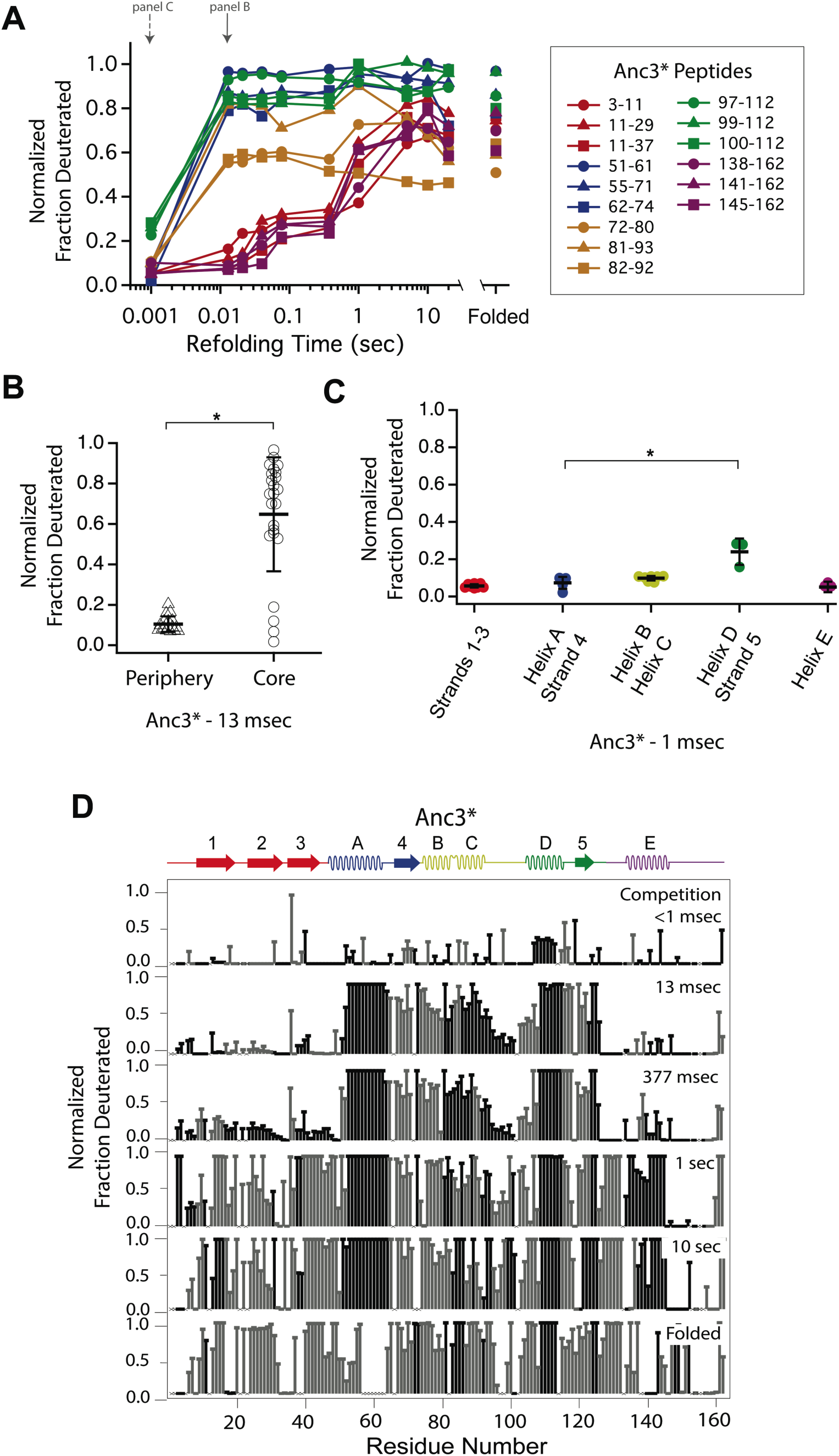
Determination of the folding pathway of Anc3* by HX-MS. **A)** Protection of representative peptides from Anc3* at various refolding times. Peptides are colored according to their corresponding structural element. The solid arrow indicates the refolding time point analyzed in panel B. The dotted arrow indicates the refolding time point analyzed in panel C. **B)** Protection of peptides mapping to the core region (I_core_) or the periphery region of Anc3* at 13 msec after refolding. Bars represent the mean and standard deviation of each data set. *p < 0.0001 (Welch’s unpaired T-test) **C)** Protection of peptides of Anc3* mapping to distinct secondary structural elements at 1 msec after refolding. Bars represent the mean and standard deviation of each data set. *p = 0.0419 (Welch’s unpaired T-test). **D)** Residue-resolved folding pathway of Anc3* at representative refolding time points. Data points in black indicate residues that are site-resolved. Data points in grey indicate residues in regions with less peptide coverage and are thus not site-resolved with the neighboring residues. Residues where site-resolved protection could not be determined due to insufficient peptide coverage is denoted with a “x”.

**Figure S4.**
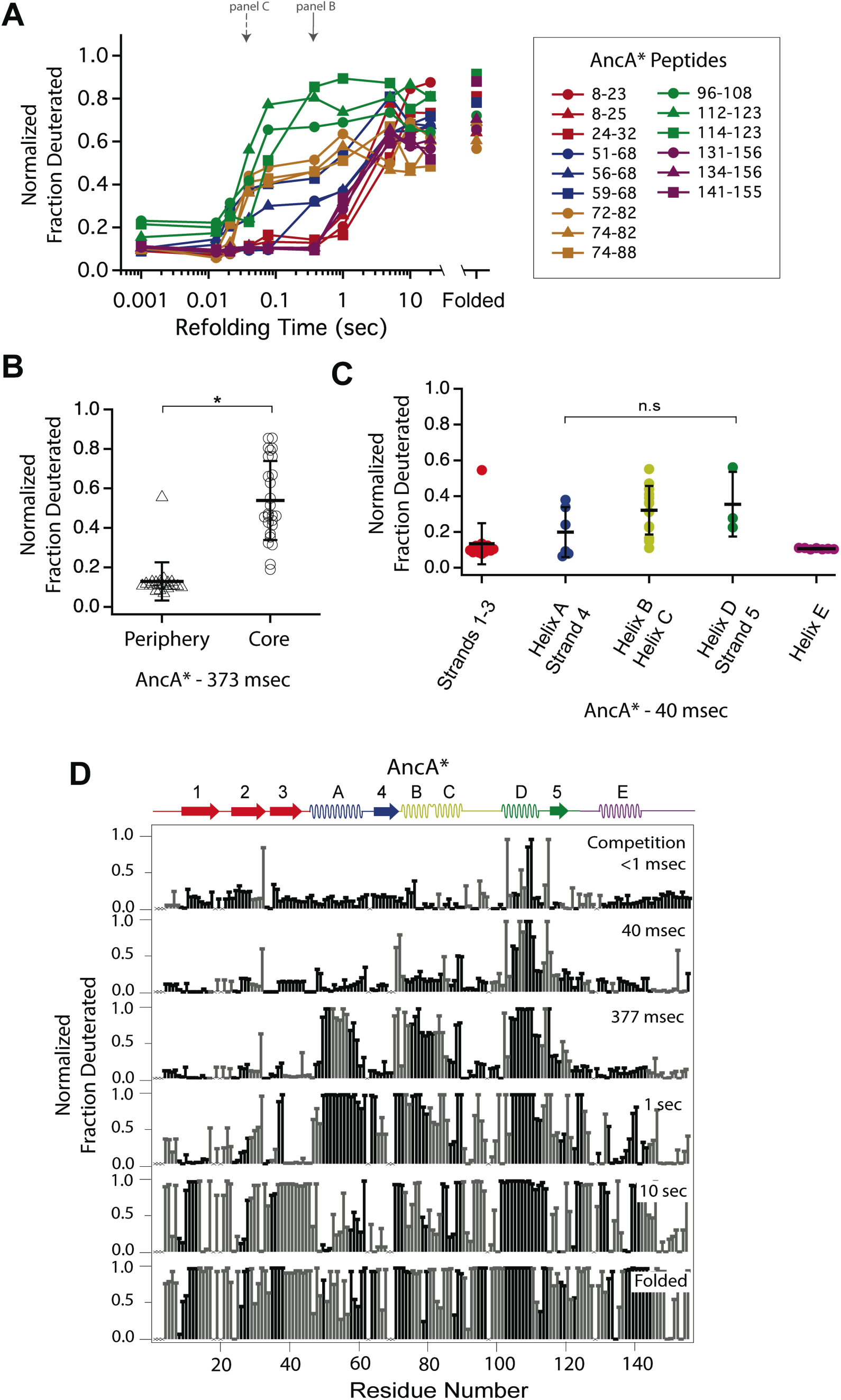
Determination of the folding pathway of AncA* by HX-MS. **A)** Protection of representative peptides from AncA* at various refolding times. Peptides are colored according to their corresponding structural element. The solid arrow indicates the refolding time point analyzed in panel B. The dotted arrow indicates the refolding time point analyzed in panel C. **B)** Protection of peptides mapping to the core region (I_core_) or the periphery region of AncA* at 373 msec after refolding. Bars represent the mean and standard deviation of each data set. *p < 0.0001 (Welch’s unpaired T-test) **C)** Protection of peptides of AncA* mapping to distinct secondary structural elements at 40 msec after refolding. Bars represent the mean and standard deviation of each data set. p = 0.275 (n.s. = not significant, Welch’s unpaired T-test). **D)** Residue-resolved folding pathway of AncA* at representative refolding time points. Data points in black indicate residues that are site-resolved. Data points in grey indicate residues in regions with less peptide coverage and are thus not site-resolved with the neighboring residues. Residues where site-resolved protection could not be determined due to insufficient peptide coverage is denoted with a “x”.

**Figure S5.**
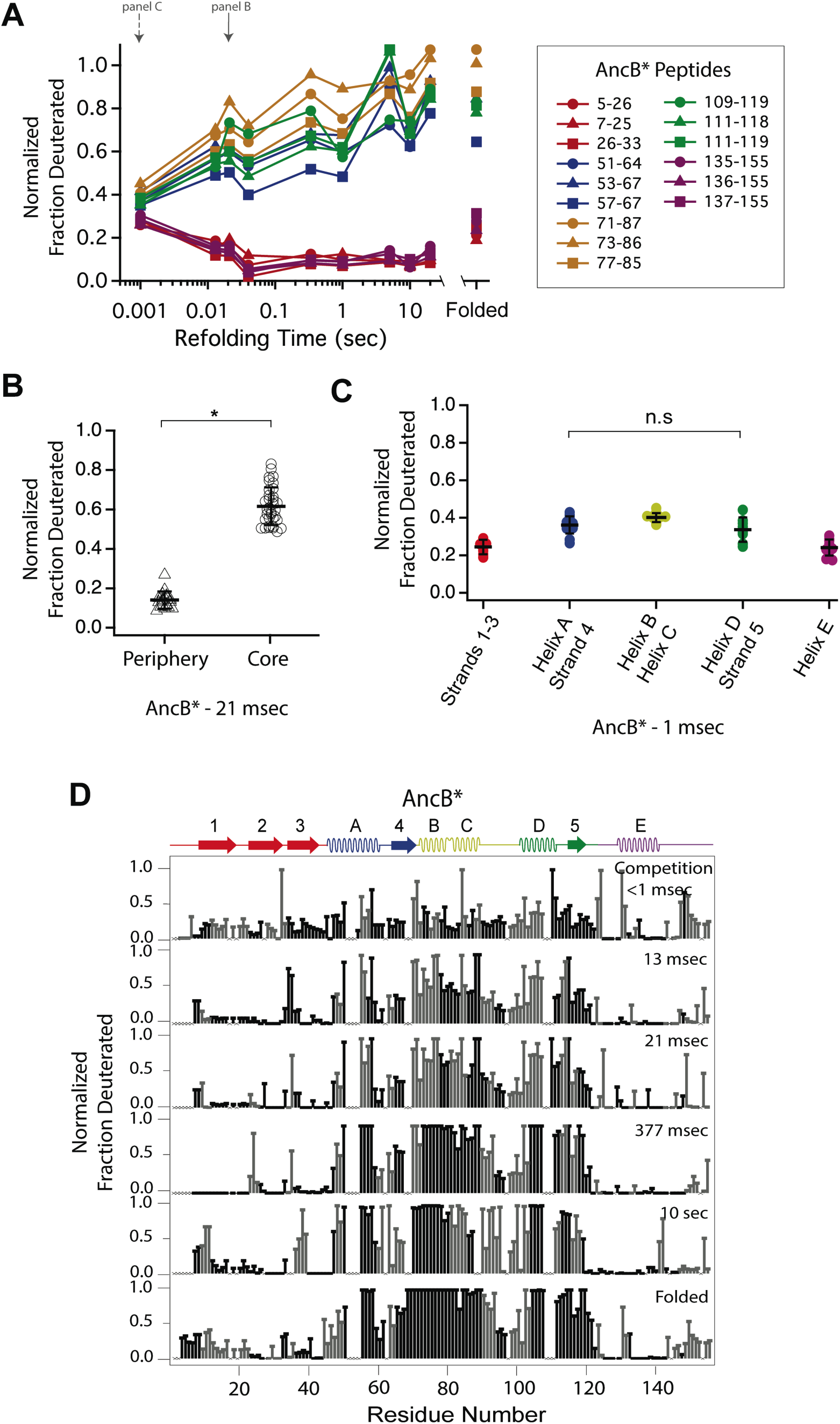
Determination of the folding pathway of AncB* by HX-MS. **A)** Protection of representative peptides from AncB* at various refolding times. Peptides are colored according to their corresponding structural element. The solid arrow indicates the refolding time point analyzed in panel B. The dotted arrow indicates the refolding time point analyzed in panel C. **B)** Protection of peptides mapping to the core region (I_core_) or the periphery region of AncB* at 21 msec after refolding. Bars represent the mean and standard deviation of each data set. *p < 0.0001 (Welch’s unpaired T-test) **C)** Protection of peptides of AncB* mapping to distinct secondary structural elements at 1 msec after refolding. Bars represent the mean and standard deviation of each data set. p = 0.353 (n.s. = not significant, Welch’s unpaired T-test). **D)** Residue-resolved folding pathway of AncB* at representative refolding time points. Data points in black indicate residues that are site-resolved. Data points in grey indicate residues in regions with less peptide coverage and are thus not site-resolved with the neighboring residues. Residues where site-resolved protection could not be determined due to insufficient peptide coverage is denoted with a “x”.

**Figure S6.**
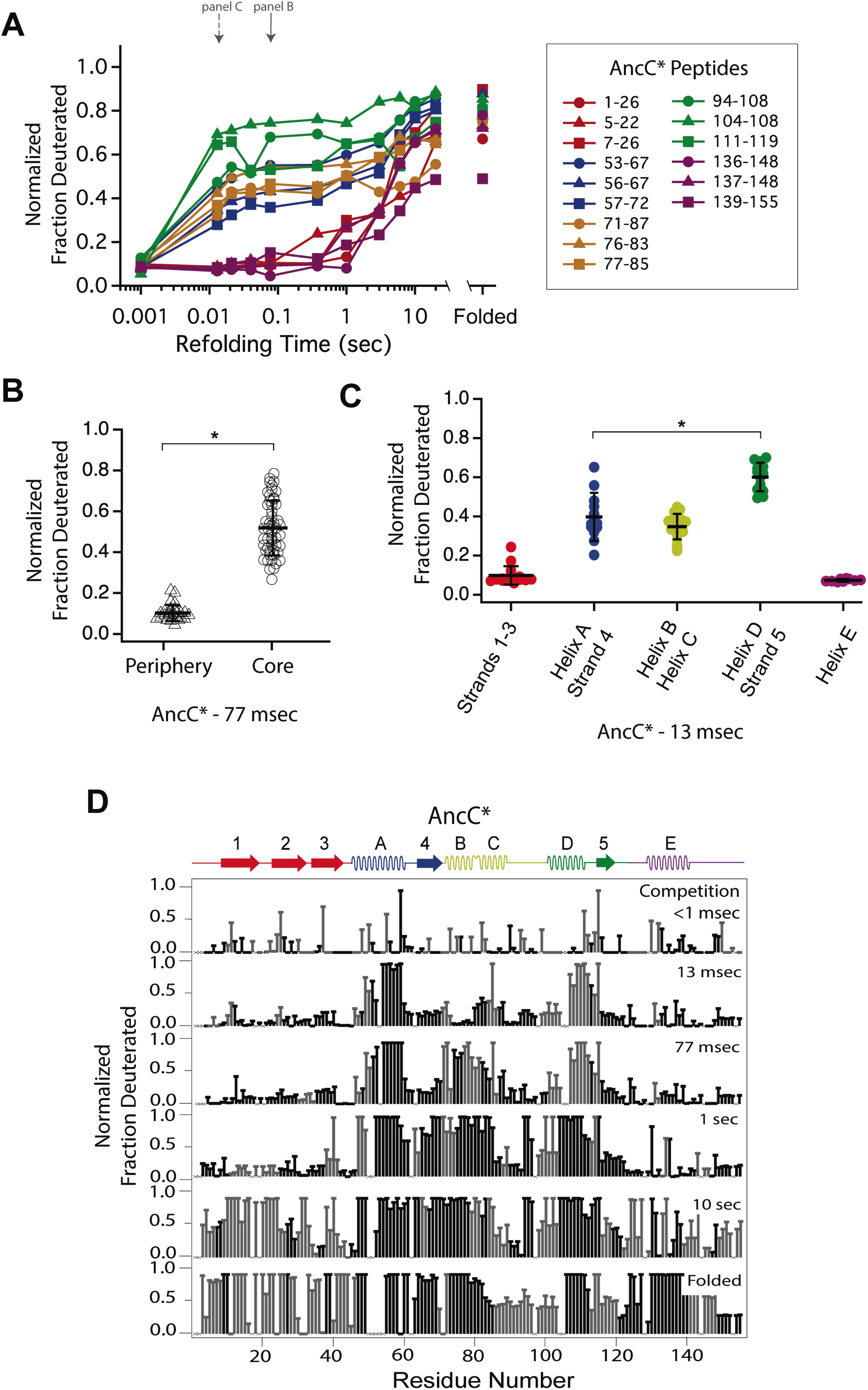
Determination of the folding pathway of AncC* by HX-MS. **A)** Protection of representative peptides from AncC* at various refolding times. Peptides are colored according to their corresponding structural element. The solid arrow indicates the refolding time point analyzed in panel B. The dotted arrow indicates the refolding time point analyzed in panel C. **B)** Protection of peptides mapping to the core region (I_core_) or the periphery region of AncC* at 77 msec after refolding. Bars represent the mean and standard deviation of each data set. *p < 0.0001 (Welch’s unpaired T-test) **C)** Protection of peptides of AncC* mapping to distinct secondary structural elements at 13 msec after refolding. Bars represent the mean and standard deviation of each data set. *p<0.0001 (Welch’s unpaired T-test). **D)** Residue-resolved folding pathway of AncC* at representative refolding time points. Data points in black indicate residues that are site-resolved. Data points in grey indicate residues in regions with less peptide coverage and are thus not site-resolved with the neighboring residues. Residues where site-resolved protection could not be determined due to insufficient peptide coverage is denoted with a “x”.

**Figure S7.**
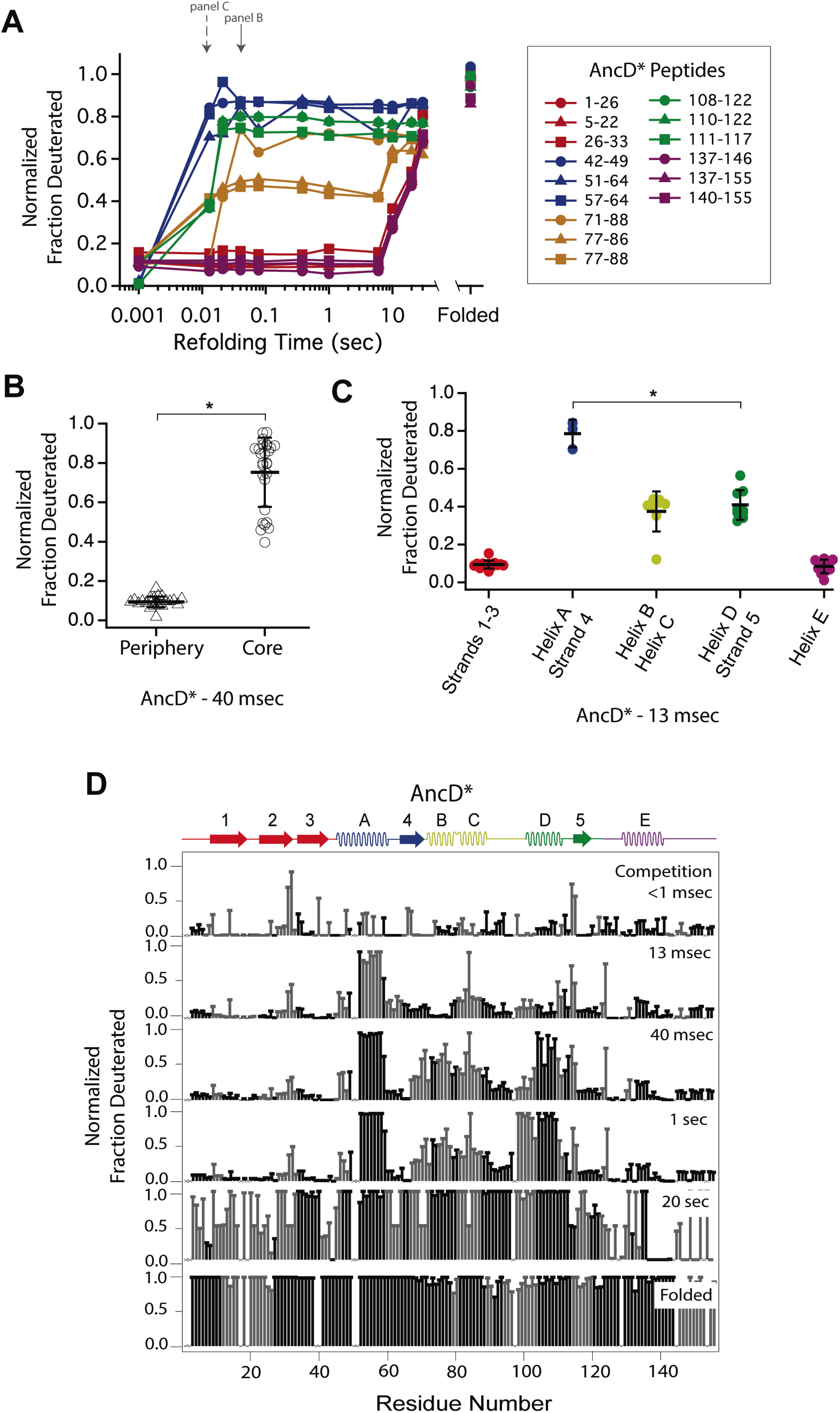
Determination of the folding pathway of AncD* by HX-MS. **A)** Protection of representative peptides from AncD* at various refolding times. Peptides are colored according to their corresponding structural element. The solid arrow indicates the refolding time point analyzed in panel B. The dotted arrow indicates the refolding time point analyzed in panel C. **B)** Protection of peptides mapping to the core region (I_core_) or the periphery region of AncD* at 40 msec after refolding. Bars represent the mean and standard deviation of each data set. *p < 0.0001 (Welch’s unpaired T-test) **C)** Protection of peptides of AncD* mapping to distinct secondary structural elements at 13 msec after refolding. Bars represent the mean and standard deviation of each data set. *p = 0.021 (Welch’s unpaired T-test). **D)** Residue-resolved folding pathway of AncD* at representative refolding time points. Data points in black indicate residues that are site-resolved. Data points in grey indicate residues in regions with less peptide coverage and are thus not site-resolved with the neighboring residues. Residues where site-resolved protection could not be determined due to insufficient peptide coverage is denoted with a “x”.

**Table S1.** Comparison of intrinsic helicity and AABUF across RNase H variants.

|  | ttRNH* |  | Anc3* |  | Anc2* |  | Anc1* |  |
| --- | --- | --- | --- | --- | --- | --- | --- | --- |
|  | AABUF (Å<sup>2</sup>)<sup>†</sup> | Helicity (%)<sup>‡</sup> | AABUF (Å<sup>2</sup>) | Helicity (%) | AABUF (Å<sup>2</sup>) | Helicity (%) | AABUF (Å<sup>2</sup>) | Helicity (%) |
| Helix A | 122.35 | **3.61** | 122.35 | **3.56** | 124.57 | **1.74** | 124.57 | **1.82** |
| Helix D | 135.97 | **8.55** | 130.04 | **5.11** | 136.91 | **17.57** | 136.91 | **20.28** |
| Ratio | 0.90 | **0.42** | 0.94 | **0.70** | 0.91 | **0.10** | 0.91 | **0.09** |
| Log2(Ratio) | -0.15 | **-1.24** | -0.09 | **-0.52** | -0.14 | **-3.34** | -0.14 | **-3.48** |

|  | AncA* |  | AncB* |  | AncC* |  | AncD* |  |
| --- | --- | --- | --- | --- | --- | --- | --- | --- |
|  | AABUF (Å<sup>2</sup>) | Helicity (%) | AABUF (Å<sup>2</sup>) | Helicity (%) | AABUF (Å<sup>2</sup>) | Helicity (%) | AABUF (Å<sup>2</sup>) | Helicity (%) |
| Helix A | 124.57 | **1.92** | 126.35 | **3.22** | 128.56 | **5.34** | 128.62 | **10.04** |
| Helix D | 130.36 | **2.59** | 131.49 | **4.63** | 136.17 | **4.05** | 129.95 | **2.48** |
| Ratio | 0.96 | **0.74** | 0.96 | **0.69** | 0.94 | **1.32** | 0.99 | **4.05** |
| Log2(Ratio) | -0.07 | **-0.43** | -0.06 | **-0.53** | -0.08 | **0.40** | -0.01 | **2.02** |

|  | ecRNH* |  | ecRNH* A55G |  | ecRNH* D108L |  |
| --- | --- | --- | --- | --- | --- | --- |
|  | AABUF (Å<sup>2</sup>) | Helicity (%) | AABUF (Å<sup>2</sup>) | Helicity (%) | AABUF (Å<sup>2</sup>) | Helicity (%) |
| Helix A | 128.62 | **9.66** | 127.23 | **3.01** | 128.62 | **9.66** |
| Helix D | 129.95 | **2.47** | 129.95 | **2.47** | 134.37 | **20.28** |
| Ratio | 0.99 | **3.92** | 0.98 | **1.22** | 0.96 | **0.48** |
| Log2(Ratio) | -0.01 | **1.97** | -0.03 | **0.29** | -0.06 | **-1.07** |

<sup>†</sup> AABUF values are the average across each helix as predicted using values from Rose, et al. (1985).
<sup>‡</sup> Helicity values are the average across each helix as predicted using values from Muñoz and Serrano (1994).

## References

1. Dill, K. A. & MacCallum, J. L. The Protein-Folding Problem, 50 Years On. Science 338, 1042–1046 (2012).

2. Baldwin, R. L. Intermediates in Protein Folding Reactions and the Mechanism of Protein Folding. Annu. Rev. Biochem. 44, 453–475 (1975).

3. Karamanos, T. K. et al. A population shift between sparsely-populated folding intermediates determines amyloidogenicity. J. Am. Chem. Soc. 138, 6271–6280 (2016).

4. Chiti, F. & Dobson, C. M. Protein Misfolding, Amyloid Formation, and Human Disease: A Summary of Progress Over the Last Decade. Annu. Rev. Biochem. 86, 27–68 (2017).

5. Ahn, M. et al. The Significance of the Location of Mutations for the Native-State Dynamics of Human Lysozyme. Biophys. J. 111, 2358–2367 (2016).

6. Hu, W. et al. Stepwise protein folding at near amino acid resolution by hydrogen exchange and mass spectrometry. Proc. Natl. Acad. Sci. 110, 7684–9 (2013).

7. Walters, B. T., Ricciuti, A., Mayne, L. & Englander, S. W. Minimizing Back Exchange in the Hydrogen Exchange-Mass Spectrometry Experiment. J. Am. Soc. Mass Spectrom. 23, 2132–2139 (2012).

8. Mayne, L. et al. Many Overlapping Peptides for Protein Hydrogen Exchange Experiments by the Fragment Separation-Mass Spectrometry Method. J. Am. Soc. Mass Spectrom. 22, 1898–1905 (2011).

9. Aghera, N. & Udgaonkar, J. B. Stepwise Assembly of β-Sheet Structure during the Folding of an SH3 Domain Revealed by a Pulsed Hydrogen Exchange Mass Spectrometry Study. Biochemistry 56, 3754–3769 (2017).

10. Vahidi, S., Stocks, B. B., Liaghati-Mobarhan, Y. & Konermann, L. Submillisecond Protein Folding Events Monitored by Rapid Mixing and Mass Spectrometry-Based Oxidative Labeling. Anal. Chem. 85, 8618–8625 (2013).

11. Khanal, A., Pan, Y., Brown, L. S. & Konermann, L. Pulsed hydrogen/deuterium exchange mass spectrometry for time-resolved membrane protein folding studies. J. Mass Spectrom. 47, 1620–1626 (2012).

12. Raschke, T. M. & Marqusee, S. The kinetic folding intermediate of ribonuclease H resembles the acid molten globule and partially unfolded molecules detected under native conditions. Nat. Struct. Biol. 4, 298–304 (1997).

13. Raschke, T. M., Kho, J. & Marqusee, S. Confirmation of the hierarchical folding of RNase H : a protein engineering study. Nat. Struct. Biol. 6, 825–831 (1999).

14. Cecconi, C., Shank, E. A., Bustamante, C. & Marqusee, S. Direct observation of the three-state folding of a single protein molecule. Science 309, 2057–2060 (2005).

15. Rosen, L. E., Connell, K. B. & Marqusee, S. Evidence for close side-chain packing in an early protein folding intermediate previously assumed to be a molten globule. Proc. Natl. Acad. Sci. 111, 14746–14751 (2014).

16. Rosen, L. E., Kathuria, S. V., Matthews, C. R., Bilsel, O. & Marqusee, S. Non-Native Structure Appears in Microseconds during the Folding of E. coli RNase H. J. Mol. Biol. 427, 443–453 (2015).

17. Chamberlain, A. K., Handel, T. M. & Marqusee, S. Detection of rare partially folded molecules in equilibrium with the native conformation of RNase H. Nat. Struct. Biol. 3, 782–7 (1996).

18. Kern, G., Handel, T. M. & Marqusee, S. Characterization of a folding intermediate from HIV-1 ribonuclease H. Protein Sci. 7, 2164–2174 (1998).

19. Hollien, J. & Marqusee, S. Comparison of the folding processes of T. thermophilus and E. coli ribonucleases H. J. Mol. Biol. 316, 327–340 (2002).

20. Ratcliff, K., Corn, J. & Marqusee, S. Structure, stability, and folding of ribonuclease H1 from the moderately thermophilic Chlorobium tepidum: comparison with thermophilic and mesophilic homologues. Biochemistry 48, 5890–5898 (2009).

21. Harms, M. J. & Thornton, J. W. Evolutionary biochemistry: revealing the historical and physical causes of protein properties. Nat. Rev. Genet. 14, 559–571 (2013).

22. Wheeler, L. C., Lim, S. A., Marqusee, S. & Harms, M. J. The thermostability and specificity of ancient proteins. Curr. Opin. Struct. Biol. 38, 37–43 (2016).

23. Starr, T. N., Picton, L. K. & Thornton, J. W. Alternative evolutionary histories in the sequence space of an ancient protein. Nature 549, 409–413 (2017).

24. Gaucher, E. A., Govindarajan, S. & Ganesh, O. K. Palaeotemperature trend for Precambrian life inferred from resurrected proteins. Nature 451, 704–707 (2008).

25. Hobbs, J. K. et al. On the origin and evolution of thermophily: reconstruction of functional precambrian enzymes from ancestors of Bacillus. Mol. Biol. Evol. 29, 825–835 (2012).

26. Perez-Jimenez, R. et al. Single-molecule paleoenzymology probes the chemistry of resurrected enzymes. Nat. Struct. Mol. Biol. 18, 592–596 (2011).

27. Risso, V. A., Gavira, J. A., Mejia-Carmona, D. F., Gaucher, E. A. & Sanchez-Ruiz, J. M. Hyperstability and Substrate Promiscuity in Laboratory Resurrections of Precambrian β-Lactamases. J. Am. Chem. Soc. 135, 2899–2902 (2013).

28. Smock, R. G., Yadid, I., Dym, O., Clarke, J. & Tawfik, D. S. De Novo Evolutionary Emergence of a Symmetrical Protein Is Shaped by Folding Constraints. Cell 164, 476–486 (2016).

29. Akanuma, S. et al. Experimental evidence for the thermophilicity of ancestral life. Proc. Natl. Acad. Sci. 110, 11067–11072 (2013).

30. Siddiq, M. A., Hochberg, G. K. & Thornton, J. W. Evolution of protein specificity: insights from ancestral protein reconstruction. Curr. Opin. Struct. Biol. 47, 113–122 (2017).

31. Hart, K. M. et al. Thermodynamic System Drift in Protein Evolution. PLoS Biol. 12, e1001994 (2014).

32. Lim, S. A., Hart, K. M., Harms, M. J. & Marqusee, S. Evolutionary trend toward kinetic stability in the folding trajectory of RNases H. Proc. Natl. Acad. Sci. 113, 13045–13050 (2016).

33. Lim, S. A. & Marqusee, S. The burst-phase folding intermediate of ribonuclease H changes conformation over evolutionary history. Biopolymers e23086 (2017). doi:10.1002/bip.23086

34. Spudich, G. M., Miller, E. J. & Marqusee, S. Destabilization of the Escherichia coli RNase H Kinetic Intermediate: Switching Between a Two-state and Three-state Folding Mechanism. J. Mol. Biol. 335, 609–618 (2004).

35. Connell, K. B., Miller, E. J. & Marqusee, S. The folding trajectory of RNase H is dominated by its topology and not local stability: a protein engineering study of variants that fold via two-state and three-state mechanisms. J. Mol. Biol. 391, 450–460 (2009).

36. Kan, Z.-Y., Walters, B. T., Mayne, L. & Englander, S. W. Protein hydrogen exchange at residue resolution by proteolytic fragmentation mass spectrometry analysis. Proc. Natl. Acad. Sci. 110, 16438–16443 (2013).

37. Kan, Z.-Y., Mayne, L., Chetty, P. S. & Englander, S. W. ExMS: Data Analysis for HX-MS Experiments. J. Am. Soc. Mass Spectrom. 22, 1906–1915 (2011).

38. Goedken, E. R., Raschke, T. M. & Marqusee, S. Importance of the C-Terminal Helix to the Stability and Enzymatic Activity of Escherichia coli Ribonuclease H. Biochemistry 36, 7256–7263 (1997).

39. Muñoz, V. & Serrano, L. Elucidating the folding problem of helical peptides using empirical parameters. Nat. Struct. Biol. 1, 399–409 (1994).

40. Bai, Y. Protein folding pathways studied by pulsed-and native-state hydrogen exchange. Chem. Rev. 106, 1757–1768 (2006).

41. Sander, C. & Schneider, R. Database of homology-derived protein structures and the structural meaning of sequence alignment. Proteins Struct. Funct. Genet. 9, 56–68 (1991).

42. Chamberlain, A. K., Fischer, K. F., Reardon, D., Handel, T. M. & Marqusee, S. Folding of an isolated ribonuclease H core fragment. Protein Sci. 8, 2251–2257 (1999).

43. Rosen, L. E. & Marqusee, S. Autonomously Folding Protein Fragments Reveal Differences in the Energy Landscapes of Homologous RNases H. PLoS One 10, e0119640 (2015).

44. Baldwin, A. J. & Kay, L. E. NMR spectroscopy brings invisible protein states into focus. Nat. Chem. Biol. 5, 808–814 (2009).

45. Boehr, D. D., McElheny, D., Dyson, H. J. & Wright, P. E. The dynamic energy landscape of dihydrofolate reductase catalysis. Science 313, 1638–1642 (2006).

46. Kanaya, S., Katsuda-Nakai, C. & Ikehara, M. Importance of the positive charge cluster in Escherichia coli ribonuclease HI for the effective binding of the substrate. J. Biol. Chem. 266, 11621–11627 (1991).

47. Zhou, Z., Feng, H., Ghirlando, R. & Bai, Y. The high-resolution NMR structure of the early folding intermediate of the Thermus thermophilus ribonuclease H. J. Mol. Biol. 384, 531–539 (2008).

48. Rose, G. D., Geselowitz, A. R., Lesser, G. J., Lee, R. H. & Zehfus, M. H. Hydrophobicity of amino acid residues in globular proteins. Science 229, 834–838 (1985).

49. Nishimura, C., Lietzow, M. A., Dyson, H. J. & Wright, P. E. Sequence Determinants of a Protein Folding Pathway. J. Mol. Biol. 351, 383–392 (2005).

50. Nishimura, C., Dyson, H. J. & Wright, P. E. Consequences of Stabilizing the Natively Disordered F Helix for the Folding Pathway of Apomyoglobin. J. Mol. Biol. 411, 248–263 (2011).

51. Robic, S., Berger, J. M. & Marqusee, S. Contributions of folding cores to the thermostabilities of two ribonucleases H. Protein Sci. 11, 381–389 (2002).

52. Hollien, J. & Marqusee, S. Structural distribution of stability in a thermophilic enzyme. Proc. Natl. Acad. Sci. 96, 13674–13678 (1999).

53. Dabora, J. M. & Marqusee, S. Equilibrium unfolding of Escherichia coli ribonuclease H: characterization of a partially folded state. Protein Sci. 3, 1401–1408 (1994).

54. Ishikawa, K. et al. Crystal Structure of Ribonuclease H from Thermus thermophilus HB8 Refined at 2·8 Å Resolution. J. Mol. Biol. 230, 529–542 (1993).

55. Katayanagi, K. et al. Structural details of ribonuclease H from Escherichia coli as refined to an atomic resolution. J. Mol. Biol. 223, 1029–1052 (1992).

## References for Supplemental Materials

1. Hollien, J. & Marqusee, S. Comparison of the folding processes of T. thermophilus and E. coli ribonucleases H. J. Mol. Biol. 316, 327–340 (2002).

2. Lim, X.-X. et al. Epitope and Paratope Mapping Reveals Temperature-Dependent Alterations in the Dengue-Antibody Interface. Structure 25, 1391–1402.e3 (2017).

3. Muñoz, V. & Serrano, L. Elucidating the folding problem of helical peptides using empirical parameters. Nat. Struct. Biol. 1, 399–409 (1994).

4. Rose, G. D., Geselowitz, A. R., Lesser, G. J., Lee, R. H. & Zehfus, M. H. Hydrophobicity of amino acid residues in globular proteins. Science 229, 834–838 (1985).

5. Nishimura, C., Prytulla, S., Jane Dyson, H. & Wright, P. E. Conservation of folding pathways in evolutionarily distant globin sequences. Nat. Struct. Biol. 7, 679–686 (2000).

